# Prevalence and risk factors profile of seropositive *Toxoplasmosis gondii* infection among apparently immunocompetent Sudanese women

**DOI:** 10.1101/572800

**Authors:** Madinna Mustafa, Fatima Fathy, Abubaker Mirghani, Mona A. Mohamed, Mohamed S. Muneer, Abdallah E. Ahmed, Mohamed Siralkhatim Ali, Rihab A. Omer, Emmanuel E. Siddig, Nouh S. Mohamed, Amjed M. Abd Elkareem

## Abstract

**Objectives:** *Toxoplasma gondii* is an opportunistic parasite that cause a clinical manifestation known as toxoplasmosis. We investigated the prevalence and potential risk factors of *T. gondii* infection among women in Khartoum, Sudan. A sero-parasitological cross-sectional study included 100 women aging between 15-50 years old was conducted in Khartoum, Sudan between January – November 2018. Serum samples were collected and investigated for presence of anti-*T. gondii* immunoglobulins.

**Results:** Mean age of the women population included was 26.75±8.25 with a range between 15 and 50 years. Sero-prevalence of *T. gondii* antibodies was 27% (27/100) with a 95% confidence interval (CI) of 18.6 – 36.8%. Among seropositive population 81% (22/27), 15% (4/27) and 4% (1/27) were seropositive for IgG antibodies, IgM antibodies and both antibodies respectively. Age group 21-30 years old had the highest frequency of detected IgG (10/45) and IgM (3/45). Married women had the highest frequency of detected IgG or IgM, 18/79 and 3/79, respectively. Risk factors analysis showed a total of 37/100 participants were having direct contact with cats and 66/100 have a frequent raw meat consumption, neither direct cats contact nor raw meat consumption had statistically significant association with seropositivity to T. gondii (P. value =0.052 and 0.565, respectively).

## Introduction

Toxoplasmosis is an opportunistic parasitic infection caused by the parasite *Toxoplasma gondii*. Toxoplasmosis constitutes a major public health problem specially in low- and middle-income countries (LMICs) (1). More than 29% of people worldwide show serological evidence of encountering *T. gondii* infection (2). Though *T. gondii* parasite stay in a dormant stage called bradyzoites (3), the parasite transforms into an active form when the immune system becomes compromised which leads to the clinical manifestation that known as toxoplasmosis (4). Decreased immunity could be attributed to infections such as HIV (5, 6), disease e.g. fatty liver disease (7), or normal alterations in physiological response as in case of pregnancy (8–11), Congenital toxoplasmosis is another form of toxoplasmosis defined by the vertical transmission *T. gondii* tachyzoites parasite from an infected pregnant woman to a fetus through the placenta (12). Congenital toxoplasmosis occurs approximately in 1 per 1000 pregnant women (13, 14). It can cause severe damages to the fetus brain, cerebral calcification, hydrocephaly, chorioretinitis, and mental retardation (15). Some patients with congenital toxoplasmosis are likely to develop other clinical manifestations like toxoplasma encephalitis (16), lymphadenopathy (17), schizophrenia (18), ophthalmitis (19), and organ dysfunction such as liver cirrhosis (20).

In Sudan, toxoplasmosis was firstly reported in 1966 in 61% patients in Darfur state (21) In 1991 it was reported that high prevalence rate in Sudan are due to some habits like consumption of raw or partially cocked liver, viscera and meat (22). In 2013, an overall seroprevalence of 43.6% has been reported in samples donated for the central blood bank and samples investigated to issue travel cards. Higher prevalence of the disease was reported in HIV patients (75%) and women with abortion history (55.5%) in Khartoum state (23).

Early diagnosis of toxoplasmosis specially among females in child bearing age is recommended since the disease can cause miscarriage, still birth or congenital toxoplasmosis to their infants (24) when they get pregnant. Therefore, pre-diagnosis even with a first line detection method specially in the LMICs such as Sudan will help in the management of pregnant females with *T. gondii* infection. This study aimed at investigating the prevalence of *T. gondii* among Sudanese women in Khartoum state, Sudan and identification of risk factors associated with seropositive candidates.

## Materials and Methods

A sero-parasitological cross-sectional study was conducted in Sharq El-Nile hospital, Khartoum, Sudan between January – November 2018. The study included a total number of 100 females age 15 to 45 years. Patients subjected to immunosuppressive factors (pregnant women, transplant, HIV patients and other immunosuppressed) were excluded from the study. Sample collection was linked with a questionnaire guided interview. The confidentiality and anonymity of the participants were maintained throughout the research steps. Demographic information included age, educational level, marital status, occupation, and family size were collected. Other expected risk factors like contact with felines, consumption of raw meat or unwashed vegetables, drinking of unpasteurized milk, source of drinking water, rearing of rabbits and previous history of spontaneous abortion were considered as well.

## Samples collection

Two ml blood samples were collected in plain vacutainers after obtaining an informed consent from human subjects included in the study. Samples were then transported to the molecular biology lab of the National University Biomedical Research Institute – National University, Sudan. Samples were then left on bench for 30 minutes at room temperature to allow blood clotting and serum separation. Serum samples were then transferred into new plain labeled containers and stored at −20 °C till further processing.

## Serological diagnosis of toxoplasmosis

A total 100 serum samples were serologically tested for IgM and IgG against *T. gondii* antigens. The sera were subjected to Toxoplasma-specific rapid diagnostic testing (RDT), according to the manufacture’s instructions (Biopanda Reagents, Belfast, UK). Briefly, 20 µl serum were added to the pore in the cellulose strip of the test device and then allowed to migrate through the whole strip. Results were positive when obtaining two or three lines of reaction, one for the control showing the validity of the test device and second one for the positive serum sample. Three lines were obtained when both IgM and IgG detected in addition to the control line. In order to obtain a valid test result, samples were run in triplicates.

RDT seropositive samples were subjected to latex indirect agglutination test card in accordance to the manufacturer’s instructions (Rapid Labs, United Kingdom). Briefly, 20 ul serum sample was placed on the test spot next to the controls’ spots; positive and negative controls. 3 drops of the antigen provided with the test kits were added and mixed slowly with the serum followed by 15 minutes incubation at room temperature. Direct agglutination confirmatory tests were done in duplicates before tabulation of the results.

## Statistical analysis

Statistical analysis was performed using the statistical package for social sciences (SPSS version 20). Demographic data were categorized and percentages were compared using the student T-test. Chi-square test was used to test the significance of risk factors associated with seropositive patients, results considered to be statistically significant if P value < 0.05.

## Results

### Study population characteristics

Our study population consisted of 100 women aged 15 to 50 years (mean age 26.75±8.25 years). Participants were grouped according to their ages into 4 groups; 15-20, 21-30, 31-40, and 41-50 years. Most of the participants (45%) were in the 21-30 years age group. Information about participants residence, marital status, water source and contact with cats is in Table 1.

### Serological detection of *T. gondii* infection

The overall prevalence of *T. gondii* infection in the study population was (27%, 95% Confidence interval [CI] 18.6 – 36.8%) (Table 2). Results of detection were further classified based on the type of immunoglobulins present in the serum. *T. gondii* -specific IgG and IgM antibodies were detected in 22 (22%) and 4 (4%) of the seropositive participants respectively. Only one serum sample was positive for both IgG and IgM anti-*T. gondii* antibodies. Women aged 21-30 years had the highest frequency of detected IgG and IgM; 10/45 (22.2%) and 3/45 (6.7%), respectively (Table 2). When analyzing the immunoglobulins results based on the martial status of the participated women, married women had the highest frequency of detected IgG or IgM, 18/79 (22.8%) and 3/79 (3.8%) respectively. Similarly, the single woman, 1/79 (1.3%), concurrently seropositive for both IgG and IgM was married. Five of the 22 married women with positive serology for anti-*T. gondii* IgG and/or IgM antibodies also have history of spontaneous abortion. However, most married women (72.1%) were negative for the presence of anti-*T. gondii* immunoglobulins (Table 2).

### Risk factors analysis

A total of 37 (37%) participants were having direct contact with cats and 66 (66%) reported frequent raw meat consumption. However, neither direct cats contact nor raw meat consumption had statistically significant association with seropositivity to *T. gondii* (P. value = 0.052 and 0.565, respectively). Other factors related to the risk of having *T. gondii* infection including consumption of unwashed vegetables, drinking of unpasteurized milk, owing of rabbits and, the source of drinking water was described in table 3.

Six (22.2%) seropositive women had a previous history of abortion. Four of those participants positive for *T. gondii* IgG and one positive for *T. gondii* IgM had history of two episodes of abortion while one woman who was seropositive for both IgM and IgG immunoglobulins had one previous abortion. Analysis of risk factors of having toxoplasmosis and its relation to the frequency of abortion is found in supplementary file 1.

## Discussion

This study showed 27% seroprevalence of anti-*T. gondii* antibodies among Sudanese women in Khartoum. This finding was lower than the seroprevalence of toxoplasmosis reported by Khalil et al., 2013 in which 45% of the sampled population had serological evidence of *T. gondii* infection (25). Also, our result was lower than those reported in different countries as Ghana (26), Cameroon (27) and Ethiopia (28–33). However, this seroprevalence of *T. gondii* antibodies was higher than that reported in European, North and West American continents (9, 34). Such variations could be due to different study settings in terms of multiple climatic conditions, heterogenous study population and risk factors profile such as contact with pets, hygienic practice, and feeding habit (28, 29, 31, 35, 36). The result of those who aged 21-30 years having the highest frequency of *T. gondii* infection (28.9%) was similar to previous reports (28, 29, 36) with the exception of one study conducted in Burkina Faso that showed these who aged more than 30 years were the most infected age group (35).

The results of risk factors associated with toxoplasmosis including direct contact with cats, consumption of raw meat, consumption of unwashed vegetables, drinking of unpasteurized milk, owing rabbits at home, and source of drinking water were statistically insignificant (P. values= 0.052, 0.565, 0.540, 0.706, 0.483 and 0.413, respectively). It is worth noting the tendency of association between seropositivity and direct contact with cats. In general, this study results were similar to other reports in which no significant association between these risk factors and having anti-*T. gondii* immunoglobulins (35, 37) and in contrast to previous reports that showed consumption of raw meat was significantly associated with toxoplasmosis infection (35, 38).

In this study, the history of two spontaneous abortion was statistically significant with having seropositive anti-*T. gondii* immunoglobulins (P. value 0.038). This result was higher compared to previous studies conducted among women with history of miscarriage in Mexico (38), but similar to spontaneous abortion rate in seropositive women for *T. gondii* infection reported in Pakistan (17%) (39), and lower than other studies done in Egypt (40), Iran (41), and India (42). When we linked the frequency of abortion to the presence of direct contact with cats, which was considered as a risk factor for having toxoplasmosis, the relationship was statistically significant (P. value = 0.031), similar to an association reported in Zambia (36).

## Conclusion

Toxoplasmosis is an opportunistic parasitic infection; apparently immunocompetent women can contract the infection with not a low rate. All women in child bearing age should be tested for *T. gondii* especially when they are planning to get pregnant to avoid the infection and its sequelae.

## Limitations

- Although the studied sample size is very small, it provides an insight in the situation of toxoplasmosis among Sudanese women. Therefore, a large-scale study with large sample size in required to provide accurate prevalence of this infection.
- Presence of the toxoplasmosis infection and miscarriage among women still unknown whether the exposure occurred before or after pregnancy therefore examining detection of infection before and after pregnancy and at different time intervals following pregnancy duration would significantly improve future studies.
- An accurate estimation for the toxoplasma antibodies in women serum is required in order to investigate active and past infections which can provide good follow up during pregnancy.

## Declarations

### Ethics approval and consent to participate

The study was approved by the National University Biomedical Research Institute Research Ethics Committee – National University, Sudan. Informed consent was obtained from each participant prior to enrollment in the study using written consent from adults, parents or legal custodians of children.

### Consent to publish

Not applicable

### Availability of Data and Materials

The datasets used and/or analyzed during the current study are available from the corresponding author on reasonable request.

### Authors’ contributions

MM, FF, AM, MAM and AMAE provided conceptual framework for the project, guidance for interpretation of the data, performed data analysis, NSM, EES, AEA, ROA and MSA participated in the performance of the Parasitological work. NSM, MSM and AMAE performed the statistical analysis and guidance for data interpretation. All authors read and approved the final manuscript.

## Acknowledgements

We are of great thanks for kind collaboration and assistance of the clinical staff of Sharq El-Nile hospital in Khartoum state, Sudan during patients’ reception and sampling. And also great thanks to all participants contributed to this work. We also thank Dr. Tarig B. Higazi, Professor of Biological Sciences, Ohio University Zanesville for his valuable comments.

