## Supplementary file 1 for "Prevalence and risk factors profile of seropositive *Toxoplasmosis gondii* infection among apparently immunocompetent Sudanese women"

Supplementary file 1: Analysis of risk factors of having toxoplasmosis and its relation to the frequency of abortion

| Risk factors | Frequency of Abortion | | | | | P. value |
| --- | --- | --- | --- | --- | --- | --- |
|  | 1 time | 2 times | 3 times | No history | Total |  |
| Toxoplasmosis |  |  |  |  |  | 0.038 |
| Seropositive | 1(3.7%) | 5(18.5%) | 0(0.0%) | 21(77.8%) | 27(27.0%) |  |
| Seronegative | 7(9.6%) | 2(2.7%) | 1(1.4%) | 63(86.3%) | 73(73.0%) |  |
| Age group |  |  |  |  |  |  |
| 15-20 years | 0(0.0%) | 0 (0.0%) | 0(0.0%) | 26 (100%) | 26(26%) | 0.000 |
| 21-30 years | 0(0.0%) | 1 (2.2%) | 0(0.0%) | 44(98.8%) | 45(45%) |  |
| 31-40 years | 4(20.0%) | 5(25.0%) | 1(5.0%) | 10(50.0%) | 20(20%) |  |
| 41-50 years | 4(45.0%) | 1(10.0%) | 0(0.0%) | 4(45.0%) | 9(9%) |  |
| Direct contact with cats |  |  |  |  |  |  |
| Yes | 0(0.0%) | 5(13.5%) | 0(0.0%) | 32(86.5%) | 37(37%) | 0.031 |
| No | 8(12.7%) | 2(3.2%) | 1(1.6%) | 52(82.5%) | 63(63%) |  |
| Consumption of raw meat |  |  |  |  |  |  |
| Yes | 7(10.6%) | 5 (7.6%) | 1(1.5%) | 53(80.3%) | 66(66%) | 0.465 |
| No | 1(2.9%) | 2(5.9%) | 0(0.0%) | 31(91.2%) | 34(34%) |  |
| Consumption of raw vegetables |  |  |  |  |  |  |
| Yes | 1(10.0%) | 1(10.0%) | 0(0.0%) | 8(80.0%) | 10(10%) | 0.954 |
| No | 7(7.8%) | 6 (6.7%) | 1(1.1%) | 76(84.4%) | 90(90%) |  |
| Consumption of unpasteurized milk |  |  |  |  |  |  |
| Yes | 0(0.0%) | 0(0.0%) | 0(0.0%) | 4(100%) | 4(4%) | 0.851 |
| No | 8(8.3%) | 7(7.3%) | 1(1.0%) | 80(83.3%) | 96(96%) |  |
| Owing rabbits at home |  |  |  |  |  |  |
| Yes | 1(16.7%) | 0(0.0%) | 0(0.0%) | 5(83.3%) | 6(6%) | 0.775 |
| No | 7(7.4%) | 7(7.4%) | 1(1.1%) | 79(84.1%) | 94(94%) |  |
| Source of drinking water |  |  |  |  |  |  |
| Tap water | 8(9.4%) | 7(8.2%) | 1(1.2%) | 69(81.2%) | 85(85%) | 0.762 |
| Filtered water | 0(0.0%) | 0(0.0%) | 0(0.0%) | 14(100%) | 14(14%) |  |
| Wells water | 0(0.0%) | 0(0.0%) | 0(0.0%) | 1(100%) | 1(1%) |  |
