## Supplementary material for "Prevalence and risk factors profile of seropositive *Toxoplasmosis gondii* infection among apparently immunocompetent Sudanese women": Table 1

Table 1: Characteristics of the study participants

| Participants Characteristics | Positive  N (%) | Negative  N (%) | Total  Out of 100 (%) | P. value |
| --- | --- | --- | --- | --- |
| Age groups |  |  |  |  |
| 15-20 | 3(11.5%) | 23(88.5%) | 26(26%) | 0.153 |
| 21-30 | 13(28.9%) | 32(71.1%) | 45(45%) |  |
| 31-40 | 7(25.9%) | 20(74.1%) | 27(27%) |  |
| 41-50 | 4(30.8%) | 9(69.2%) | 13(13%) |  |
| Marital Status |  |  |  |  |
| Married | 22(27.8%) | 57(72.2%) | 79(79%) | 0.855 |
| Single | 4(26.7%) | 11(73.3%) | 15(15%) |  |
| Divorced | 0(0.0%) | 2(100%) | 2(2%) |  |
| Widow | 1(25.0%) | 3(75.0%) | 4(4%) |  |
| Residence |  |  |  |  |
| Al Fayha | 8(33.3%) | 16(66.7%) | 24(24%) | 0.491 |
| Al Gadesia | 0(0.0%) | 3(100%) | 3(3%) |  |
| Gereaf sharq | 10(34.5%) | 19(65.5%) | 29(29%) |  |
| Haj Yousef | 7(21.2%) | 26(78.8%) | 33(33%) |  |
| Soba Sharq | 2(18.2%) | 9(81.8%) | 11(11%) |  |
| Direct contact with cats |  |  |  |  |
| Yes | 14(37.8%) | 23(62.2%) | 37(37%) | 0.052 |
| No | 13(20.6%) | 50(79.4%) | 63(63%) |  |
| Consumption of raw meat |  |  |  |  |
| Yes | 18(27.3%) | 48(72.7%) | 66(66%) | 0.565 |
| No | 9(26.5%) | 25(73.4%) | 34(34%) |  |
| Consumption of raw vegetables |  |  |  |  |
| Yes | 3(30.0%) | 7(70.0%) | 10(10%) | 0.540 |
| No | 24(26.7%) | 66(73.3%) | 90(90%) |  |
| Consumption of unpasteurized Milk |  |  |  |  |
| Yes | 1(25.0%) | 3(75.5%) | 4(4%) | 0.706 |
| No | 26(27.1%) | 70(72.9%) | 96(96%) |  |
| Owing rabbits at home |  |  |  |  |
| Yes | 1(16.7%) | 5(83.3%) | 6(6%) | 0.483 |
| No | 26(27.7%) | 68(72.3%) | 94(94%) |  |
| Source of Drinking water |  |  |  |  |
| Tap water | 25(29.4%) | 60(70.6%) | 85(85%) | 0.413 |
| Filtered water | 2(14.3%) | 12(85.7%) | 14(14%) |  |
| Wells water | 0(0.0%) | 1(100%) | 1(1%) |  |
| History of Abortion |  |  |  |  |
| Yes | 6(37.5%) | 10(62.5%) | 16(16%) | 0.230 |
| No | 21(25.0%) | 63(75.0%) | 84(84%) |  |
