## Supplementary material for "Prevalence and risk factors profile of seropositive *Toxoplasmosis gondii* infection among apparently immunocompetent Sudanese women": Table 2

Table 2: Results of serological diagnosis among the study participants

|  | Immunoglobulins | | | | Total | P. value |
| --- | --- | --- | --- | --- | --- | --- |
|  | IgG | IgM | IgG and IgM | Negative |  |  |
| Age groups |  |  |  |  |  |  |
| 15-20 years | 3(11.5%) | 0(0.0%) | 0(0.0%) | 23(88.5%) | 26(26%) | 0.063 |
| 21-30 years | 10(22.2%) | 3(6.7%) | 0(0.0%) | 32(71.1%) | 45(45%) |  |
| 31-40 years | 6(30.0%) | 1(5.0%) | 0(0.0%) | 13(65.0%) | 20(20%) |  |
| 41-50 years | 3(33.3%) | 0(0.0%) | 1(11.1%) | 5(55.6%) | 9(9%) |  |
| Marital Status |  |  |  |  |  |  |
| Married | 18 (22.8%) | 3 (3.8%) | 1 (1.3%) | 57(72.1%) | 79 (97.0%) | 0.997 |
| Single | 3 (20.0%) | 1 (6.7%) | 0 (0.0%) | 11 (73.3%) | 15 (15.0%) |  |
| Divorced | 0 (0.0%) | 0 (0.0%) | 0 (0.0%) | 2 (100%) | 2 (2.0%) |  |
| Widow | 1 (25.0%) | 0 (0.0%) | 0 (0.0%) | 3 (75.0%) | 4 (4.0%) |  |
| frequency of abortion |  |  |  |  |  |  |
| 1 time | 0 (0.0%) | 0 (0.0%) | 1 (12.5%) | 6 (87.5%) | 8 (8.0%) | 0.009 |
| 2 times | 4 (57.1%) | 1 (14.3%) | 0 (0.0%) | 2 (28.6%) | 7 (7.0%) |  |
| 3 times | 0 (0.0%) | 0 (0.0%) | 0 (0.0%) | 1 (100%) | 1 (1.0%) |  |
| No history of abortion | 18 (21.4%) | 3 (3.6%) | 0 (0.0%) | 61 (72.6%) | 84 (84.0%) |  |
