## Supplementary material for "Prevalence and risk factors profile of seropositive *Toxoplasmosis gondii* infection among apparently immunocompetent Sudanese women": Table 3

Table 3: Risk factors analysis with the prevalence of toxoplasmosis among the seropositive participants

| Risk Factors* | Yes | No | P. value | 95% Confidence Interval |
| --- | --- | --- | --- | --- |
| Contact with cats | 14 | 13 | 0.052 | 0.95-5.77 |
| Consumption of raw meat | 18 | 9 | 0.565 | 0.49-2.65 |
| Consumption of unwashed vegetables | 3 | 24 | 0.540 | 0.28-4.92 |
| Consumption of unpasteurized milk | 1 | 24 | 0.706 | 0.08-9.01 |
| Owing rabbits at home | 1 | 26 | 0.483 | 0.05-4.69 |
| History of abortion | 6 | 21 | 0.230 | 0.58-5.55 |

*Calculated for positive participants for toxoplasma positive serum
